# Interpretable deep learning for chromatin-informed inference of transcriptional programs driven by somatic alterations across cancers

**DOI:** 10.1101/2021.09.07.459263

**Authors:** Yifeng Tao, Xiaojun Ma, Drake Palmer, Russell Schwartz, Xinghua Lu, Hatice Ulku Osmanbeyoglu

## Abstract

Cancer is a disease of gene dysregulation, where cells acquire somatic and epigenetic alterations that drive aberrant cellular signaling. These alterations adversely impact transcriptional programs and cause profound changes in gene expression. Interpreting somatic alterations within context-specific transcriptional programs will facilitate personalized therapeutic decisions but is a monumental task. Toward this goal, we develop a partially interpretable neural network model called **C**hromatin-informed **I**nference of **T**ranscriptional **R**egulators **U**sing **S**elf-attention mechanism (CITRUS). CITRUS models the impact of somatic alterations on transcription factors and downstream transcriptional programs. Our approach employs a self-attention mechanism to model the contextual impact of somatic alterations. Furthermore, CITRUS uses a layer of hidden nodes to explicitly represent the state of transcription factors (TFs) to learn the relationships between TFs and their target genes based on TF binding motifs in the open chromatin regions of tumor samples. We apply CITRUS to genomic, transcriptomic, and epigenomic data from 17 cancer types profiled by The Cancer Genome Atlas. CITRUS predicts patient-specific TF activities and reveals transcriptional program variations between and within tumor types. We show that CITRUS yields biological insights into delineating TFs associated with somatic alterations in individual tumors. Thus, CITRUS is a promising tool for precision oncology.

## Introduction

The complex interplay between signaling inputs and transcriptional responses dictates important cellular functions. Dysregulation of this interplay leads to development and progression of disease, which has been most clearly delineated in the context of certain cancers. Cancer cells acquire somatic alterations that modify signaling and transcriptional programs, leading to profound changes in gene expression. We still lack a complete understanding of how somatic alterations affect cellular function in cancer. To begin to understand these effects, it is important to study somatic alterations within the specific transcriptional context in which they are found. Context- and patient-specific studies can be achieved with machine learning techniques, which are expected to facilitate personalized therapeutic decisions.

In the last decade, a monumental effort has been made to molecularly profile tumors by consortia, including The Cancer Genome Atlas (TCGA) and the International Cancer Genome Consortium (1,2). The multimodal datasets generated by these efforts include gene expression and somatic alterations, such as recurrent mutations and copy number variations (CNVs). The combination of genomic and transcriptomic information enables the integration of transcriptional states with upstream signaling pathways. Several methods have been developed to connect somatic alterations to a prior network or to gene expression (3–9). More recently, the Genomic Data Analysis Network generated assay for transposase-accessible chromatin with high-throughput sequencing (ATAC-seq) data for a subset of TCGA samples (~500 patients) (10). Although chromatin profiling helps uncover context-dependent and/or non-linear effects of transcription factors (TFs) on gene expression, it has not yet been incorporated into methods that connect somatic alterations to transcriptional programs across cancers. Incorporating DNA sequence information at promoter, intronic, and intergenic enhancers from ATAC-seq tumor profiles using TF motif analysis will improve the modeling of transcriptional regulation and delineate the impact of somatic alterations on transcriptional programs.

Deep learning is a powerful tool for capturing non-linear feature interactions that can explain the underlying biological phenomena. For example, attention mechanism is a deep learning method that has been widely used in computer vision and natural language processing. In contrast to traditional deep learning methods, the self-attention mechanism considers the contextual relationship of the input features and assigns attention weights to each input (11). In general, attention mechanisms improve the performance of deep learning models and increase the interpretability of the models. More recently, attention mechanisms have been applied to cancer genomics for cancer driver gene detection (12), drug response prediction (13), and base editing outcome prediction (14). For example, the genomic impact transformer (GIT) model utilizes a self-attention mechanism to encode the effects of somatic alterations in cancer and uses multi-layer perceptrons to predict differentially expressed genes (12). The attention mechanism enables GIT to select driver mutations that are likely to lead to downstream phenotypes. However, the GIT model lacks interpretability in the sense that it does not model intermediate TFs during modeling signaling from somatic alterations to gene expression programs.

Here, we present **C**hromatin-informed **I**nference of **T**ranscriptional **R**egulators **U**sing **S**elf-attention mechanism (CITRUS), a partially interpretable neural network model with encoder-decoder architecture. CITRUS links somatic alterations to transcriptional programs by modeling the statistical relationships between mutations, CNVs, gene expression, and TF-target gene information derived from ATAC-seq (**Fig. 1**). We show that CITRUS yields important biological insights into dysregulated TFs in individual tumors. Using a systematic *in silico* knock out approach, we identified key TFs associated with major somatic alterations. We believe CITRUS will assist researchers in providing actionable hypotheses for follow-up experiments and developing personalized and targeted therapeutics in a pan-cancer setting.

**Fig. 1:**
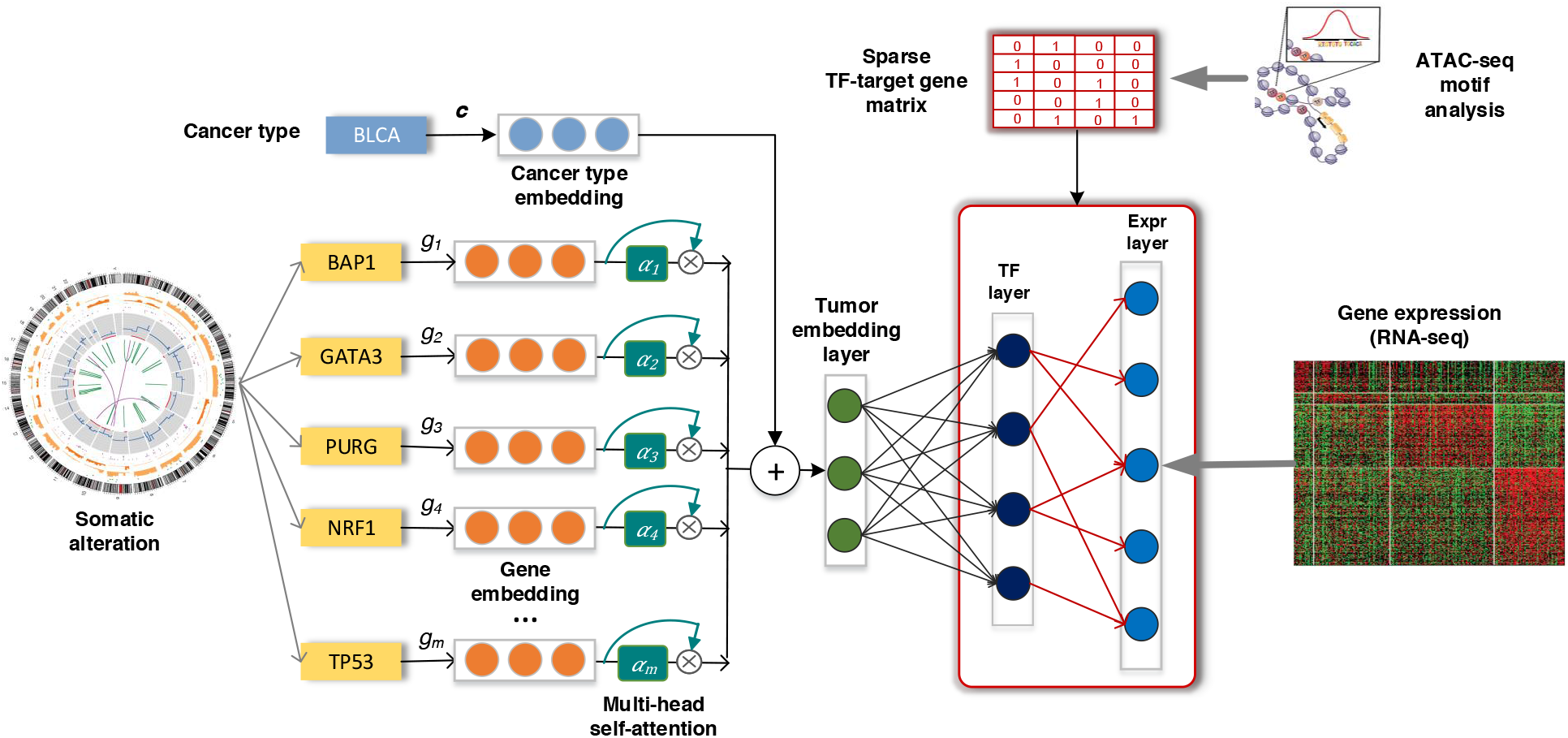
Overview of CITRUS: An attention-based model with TF-target gene priors. The input to our framework includes somatic alteration and copy number variation, assay for transposase-accessible chromatin with high-throughput sequencing (ATAC-seq), tumor expression datasets and TF recognition motifs. CITRUS takes somatic alteration and copy number variation data as input and encodes them as a tumor embedding using a self-attention mechanism. Additional cancer type information is used to stratify the confounding factor of tissue type. The middle layer further transforms the tumor embeddings into a TF layer, which represents the inferred activities of 320 TFs. Finally, gene expression levels are predicted from the TF activities through a TF-target gene priors constrained sparse layer based on ATAC-seq.

## Material & Methods

### Data pre-processing

We downloaded the batch normalized RNA-Seq expression levels quantified by RNA-Seq by Expectation Maximization (RSEM) from the Genomic Data Commons (GDC) portal (https://gdc.cancer.gov/about-data/publications/pancanatlas. We log2-transformed RSEM values and identified the 2,500 most variable genes across samples within a cancer type. Then, we took the union of the identified genes across cancer types. The final gene set included 5541 genes.

We obtained processed gene-level somatic alterations for each cancer patient from Cai et al. (4). Genes with non-synonymous mutations, small insert/deletion, or somatic copy number alteration (deletion or amplification) were given a value of 1, and otherwise were given a value of 0. We removed genes that were not present in at least 4% of samples for each cancer type.

We downloaded the ATAC-seq pan-cancer dataset from the GDC portal (https://gdc.cancer.gov/about-data/publications/ATACseq-AWG) (10). Using the MEME (15) curated Cis-BP (16) TF-binding motif reference, we scanned the pan-cancer ATAC-seq peak atlas with FIMO (17) to find peaks likely to contain each motif (*P* < 10^−5^). The final set contained 320 motifs. We associated each peak with its nearest gene in the human genome using the ChIPpeakAnno package (18). ATAC-seq peaks located in the body of the transcription unit, 100 kb upstream of the transcription start site (TSS), and 100 kb downstream of the 3’ end were assigned to the associated gene. TF-binding site identification was used to convert the assigned ATAC peaks for each gene to a feature vector of binding signals by assigning the maximum score of each motif across all peaks to a gene. Then, we created a matrix **C** ∈ {0,1}^kxl^ that defines a candidate set of associations between TFs and target genes. C_i,j_ = 1 when there is a connection from TF *j* to the gene/RNA *i* (red lines connecting the TF layer and target gene expression (Exp) layer in **Fig. 1**).

### CITRUS model

CITRUS is a framework for modeling impact of somatic alterations on transcriptional programs. **Fig. 1** shows the model architecture with an overall encoder and decoder structure. Somatic gene alteration inputs with more than 20K dimensions were encoded into a compressed representation as tumor embedding and then decode to a large dimension data of gene expression. This allows the model to capture key features of the high dimension inputs and reduce the data noise as well.

We designed a self-attention mechanism which assigned importance weights to input features (somatic alterations) through the model training. Formally, given a specific tumor *t*, with the cancer type *s*, we have a set of somatic alterations in the tumor 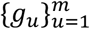 where *m* is number of mutant genes. The encoder module first maps each gene *g* (it is *g*_*u*_ here, but we omit the subscript for notation simplicity) into its corresponding gene vector ***e***_*g*_. Then, the encoder utilizes the multi-head self-attention mechanism to calculate the weighted sum of both the gene embeddings and the cancer type embedding:

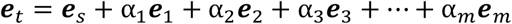

The self-attention mechanism takes the gene embeddings of all mutated/altered genes as an input and outputs the attention weights 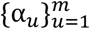 through a sub-neural network. The attention mechanism captures the context of co-existing somatic alterations and their complex interactions, which is lost in simpler models. Interested readers can find the mathematical details of self-attention mechanisms in the cited reference (12).

The decoder first infers the TF activities from the encoded tumor embedding ***e***_*t*_:

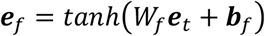

We used tanh activation instead of ReLU operation, which is more widely used in deep learning, because it has similar performance to that of ReLU in our model and generates more biologically meaningful results (e.g., distribution of TFs ***e***_*f*_). Finally, CITRUS predicts cancer type-specific mRNA expression from TF activities:

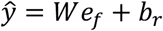

where *W* corresponds to the sparse TF-target gene matrix constrained by the prior *C* ∈ {0,1}^*k×l*^. More specifically, to integrate priors into our model, *W* shares the same shape with prior *C*, and *W*_*i,j*_ is allowed to be nonzero only when *C*_*i,j*_ = 1, and W_i,j_ is constrained to be non-negative value. We use mean square loss function as: 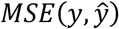

One might use other common approaches to integrate prior **C** into the **W**, i.e., by applying a Gaussian prior to *W*, which is equivalent to adding an additional penalty to the loss function 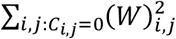. However, this “soft” constraint tends to generate less stable TF layers across different runs of training compared to the “hard” constraints shown in our model.

To prevent overfitting and to increase robustness to noise, we introduced additional dropout operations with a dropout rate of 0.2 after the input layer, activated tumor embedding layer, and activated TF layer.

### Training and evaluation

We implemented CITRUS through the PyTorch package (https://pytorch.org/), and training was performed using the Adam optimizer with default parameters except for the learning rate^15^ and weight decay. We set the learning rate to 1 × 10^−3^ and the weight decay to 1 × 10^−5^. We used early stopping with patience of 30 steps to stop training.

For statistical evaluation, we computed the mean Spearman correlation (ρ) between predicted and measured gene expression profiles for each tumor type. We split datasets into training (40%), validation (20%), and testing (20%) sets. For CITRUS, we utilized the training and validation sets to tune hyperparameters, such as the learning rate and training steps, and then evaluated these parameters on the testing set. For affinity regression (see below), we separated datasets by cancer type and conducted 5-fold cross-validation to tune hyperparameters in the training and validation sets. Then, we applied the trained model with selected hyperparameters to the testing set for performance evaluation. To increase the stability of inferred TF activity analysis, we assembled multiple CITRUS models trained with different random initialization state and integrated the TF layer based on the average of 10 trials.

#### Parameter selection

CITRUS includes more than 10 hyperparameters that are described in the following paragraphs. These hyperparameters were tuned for optimal performance in the validation set. Ideally, hyperparameter optimization is performed using a grid search of all parameters. However, this is not practical due to the tremendous computational cost. For example, three options for each parameter leads to 3^10^ possible combinations for just 10 parameters. In addition, we guide the performance metric by k-fold cross-validation, and the total experiments necessary would be 5×3^10^ (k=5). Therefore, our hyperparameter tuning strategy combined automatic and manual tuning. First, we created empirical settings for each parameter and randomly selected a set of parameters from 100 combinations. We utilized the best-performing settings to narrow down the preliminary decisions and correlation among parameters. Then, we tuned parameters independently or in sub-groups manually or by grid search.

#### Model robustness

The learning rate is perhaps the most important hyperparameter in neural network training. We first tested the learning rate in a range of settings [10^−5^, 10^−4^, 10^−3^, 10^−2^ ...], starting with the lowest setting and progressing to larger values until validation loss started to diverge. We found that if the learning rate was too small, overfitting occurred and picked up input noise. Additionally, overfitting reduced the number of driver genes that were covered in downstream attention weight analyses. If the learning rate was too big, however, the model could not converge to an optima and yielded higher validation loss. Ultimately, we selected learning rates of 10^−3^ and 10^−4^ and applied a weight penalty (weight decay) to find an optimal combination of settings. We set the weight decay range from 10^−6^ to 10^−4^ and performed a grid search. The optimal settings for learning rate and weight decay were determined to be 10^−3^ and 10^−5^, respectively. Although large batch sizes can accelerate learning rates and training, our experiments indicated that a learning rate of 10^−3^ was the largest value that maintained validation accuracy when tested on increasing batch sizes (16, 64, 100, and 300, which is the maximum value that could run in GPU). We found that larger batch sizes tended to have slightly higher gene-wise correlation at the cost of longer training time. To balance execution time, we selected a batch size of 100. The early stopping patience setting is also related to the learning rate and batch size. Specifically, higher learning rates and larger batch sizes require smaller patience to stop training. Higher patience settings may otherwise cause overfitting. Using our selected learning rate and batch size settings, a patience of 30 was generally sufficient to maintain training without stopping too early (underfitting) due to fluctuation and without halting too far from the optima (overfitting). We validated a patience setting of 30 by comparing it with a case of overfitting. We selected the lowest loss point in the overfit training and measured how far it was from the model with early stopping. During early stages of training, the model showed an initial drop in validation performance followed by a rise. To avoid this inconsistency, we did not apply early stopping for the first 180 test steps. To test the attention mechanism, we created a mesh grid for two attention sizes (256, 128) and four attention head settings (32, 16, 8, 4). We then performed an exhaustive grid search within these settings. Based on prediction performance, we selected 256 and eight as the optimal values for attention size and attention head, respectively.

Finally, we fine-tuned our model by adjusting the dropout rate. Because we used weight decay for regularization, dropout is considered a secondary regularization for our model. In addition to hidden layer dropout, we also applied dropout to our input to reduce input noise and network redundancy and to generate a more stable hidden TF layer. We tested a sequence of five dropout rates (0.1, 0.2, 0.3, 0.4, 0.5). All dropout rate settings yielded performances above 0.9 for average sample correlation in the testing set. We determined the dropout rate optimal value (0.2) primarily based on driver gene coverage in self-attention analyses.

As we used an early stopping mechanism, we set the maximum iteration parameter to 1000. This setting ensures that the training process stops either once the patience setting is satisfied or once the maximum iterations is reached. Code testing and quick runs were performed with a maximum iteration of one.

We tested two activation functions: ‘ReLU’ and ‘tanh’. Although both activation functions performed similarly, ‘tanh’ generated more biologically meaningful results and was selected. We also tested l2, minimax, and standard normalization (scale) to normalize gene expression and found that scale normalization generated the best prediction accuracy for our model settings.

### Training the affinity regression (AR) models

AR is an algorithm for efficiently solving a regularized bilinear regression problem (19,20) and was defined in our model as follows. For a data set of *M* tumor samples profiled using RNA-seq with *N* genes, we let **Y∈***R*^*N×M*^ be the log10 gene expression profiles of tumor samples. Each column of **Y** corresponds to an RNA-seq experiment for a cancer type. We define the TF attributes of each gene in a matrix **D ∈** *R*^*N×Q*^, where each row represents a gene, and each column represents a TF vector. The TF vector indicates whether there is a binding site for the TF on each gene based on ATAC-seq data. We define the somatic alteration attributes of tumor samples as a matrix **P ∈** *R*^*M×S*^ where each row represents a tumor sample, and each column represents the somatic alteration status for the tumor sample. We set up a bilinear regression problem to learn the weight matrix **W ∈** *R*^*Q×S*^ on paired TF and somatic alteration features:

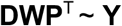

We can transform the system to an equivalent system of equations by reformulating the matrix products as Kronecker products:

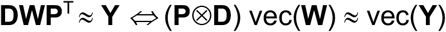

where ⊗ is a Kronecker product, and vec is a vectorizing operator that stacks a matrix and produces a vector. The result of this system is a standard (if large-scale) regression problem. Full details and a derivation of the reduced optimization problem are provided elsewhere (20).

### *In silico* knockout analysis

We implemented an *in silico* knock out approach that removes a specific somatic mutation (or copy number variation) *g* from all the tumor samples that carry it. The new somatic alteration profiles and the CITRUS-inferred TF activities generate a “wild type” corpus that does not contain the alteration *g*. In contrast, the original samples containing the alteration *g* serve as the “mutant/altered” group. We then conducted t-tests between the mutant and wild type groups to evaluate the impact of mutation *g*. This approach captures the contextual effects of mutations through the non-linear attention module of CITRUS and provides a controlled experimental environment that holds all mutations constant except for mutation *g*. For complex genotypes, the model explains TF activity across tumors. We then corrected for multiple hypotheses across models, treating inferred TF activities as separate groups of tests.

### Statistical analysis

Statistical tests were performed with the R statistical environment (4.0.2) and *Python*. For population comparisons of inferred TF activities, we performed Student’s t-tests and determined the direction of shifts by comparing the mean of the two populations. We corrected raw *P*-values for multiple hypothesis testing based on two methods: Bonferroni and FDR (BH method).

Association score between TF activity subtypes and frequent somatic alterations. For each somatic mutation or copy number variation, we calculated the *P*-value of its frequency in a cancer subtype compared to other subtypes using Fisher’s exact test. The *P*-value was further adjusted through FDR across subtypes. To identify the relative frequency of a somatic alteration in a subtype, we defined an association score, which is the product of the relative frequency direction and −log_10_FDR.

## Results

### Pan-cancer modeling of transcriptional programs

To systematically interpret somatic alterations within context-specific transcriptional programs and to identify disrupted TFs that drive tumor-specific gene expression patterns across multiple cancer types, we developed CITRUS (**Fig. 1**). CITRUS traces biological signaling from somatic alterations to signaling pathways, to TFs, and finally to target gene expression (mRNA levels). To enable this tracing, CITRUS employs an encoder-decoder architecture (**Fig. 1**). The encoder module compresses input somatic alterations into a latent vector variable called a tumor embedding. The decoder predicts TF activities from the tumor embedding and then predicts target gene expression. We used sparse TF-target gene priors based on tumor ATAC-seq data. Briefly, we started with an atlas of chromatin accessible genomic locations derived from the tumor types to be analyzed using ATAC-seq profiling data (see Methods). We then represented every gene by its feature vector of TF-binding scores, where motif information was summarized across all promoter, intronic, and intergenic chromatin accessible sites assigned to the gene (see Methods).

We applied this approach to 17 tumors from TCGA and identified key TFs associated with somatic alterations. Our dataset included samples from 17 different tumor types for which mRNA, somatic mutation, copy number variation, and ATAC-seq data were available: bladder urothelial carcinoma (BLCA, n=371), breast cancer (BRCA, n=719), cervical squamous cell carcinoma and endocervical adenocarcinoma (CESC, n=267), colorectal adenocarcinoma (COAD, n=271), esophageal carcinoma (ESCA, n=170), glioblastoma multiforme (GBM, n=143), head and neck squamous carcinoma (HNSC, n=475), kidney renal cell-clear carcinoma (KIRC, n=357), kidney renal papillary cell carcinoma (KIRP, n=272), liver hepatocellular carcinoma (LIHC, n=336), lung adenocarcinoma (LUAD, n=459), lung squamous cell carcinoma (LUSC, n=430), pheochromocytoma and paraganglioma (PCPG, n=109), prostate cancer (PRAD, n=449), stomach adenocarcinoma (STAD, n=373), thyroid carcinoma (THCA, n=216), and uterine corpus endometrial carcinoma (UCEC, n=361).

For statistical evaluation, we computed the mean Spearman correlation between predicted and measured gene expression profiles on the testing set (see Methods). CITRUS achieved significantly better performance than a regularized bilinear regression algorithm called affinity regression (AR) (20–22) that was trained independently for each cancer type. and explain gene expression across tumors in terms of somatic alteration status and presence of TF binding sites based on a pan-cancer ATAC-seq atlas (**Fig. 2A**).

**Fig. 2:**
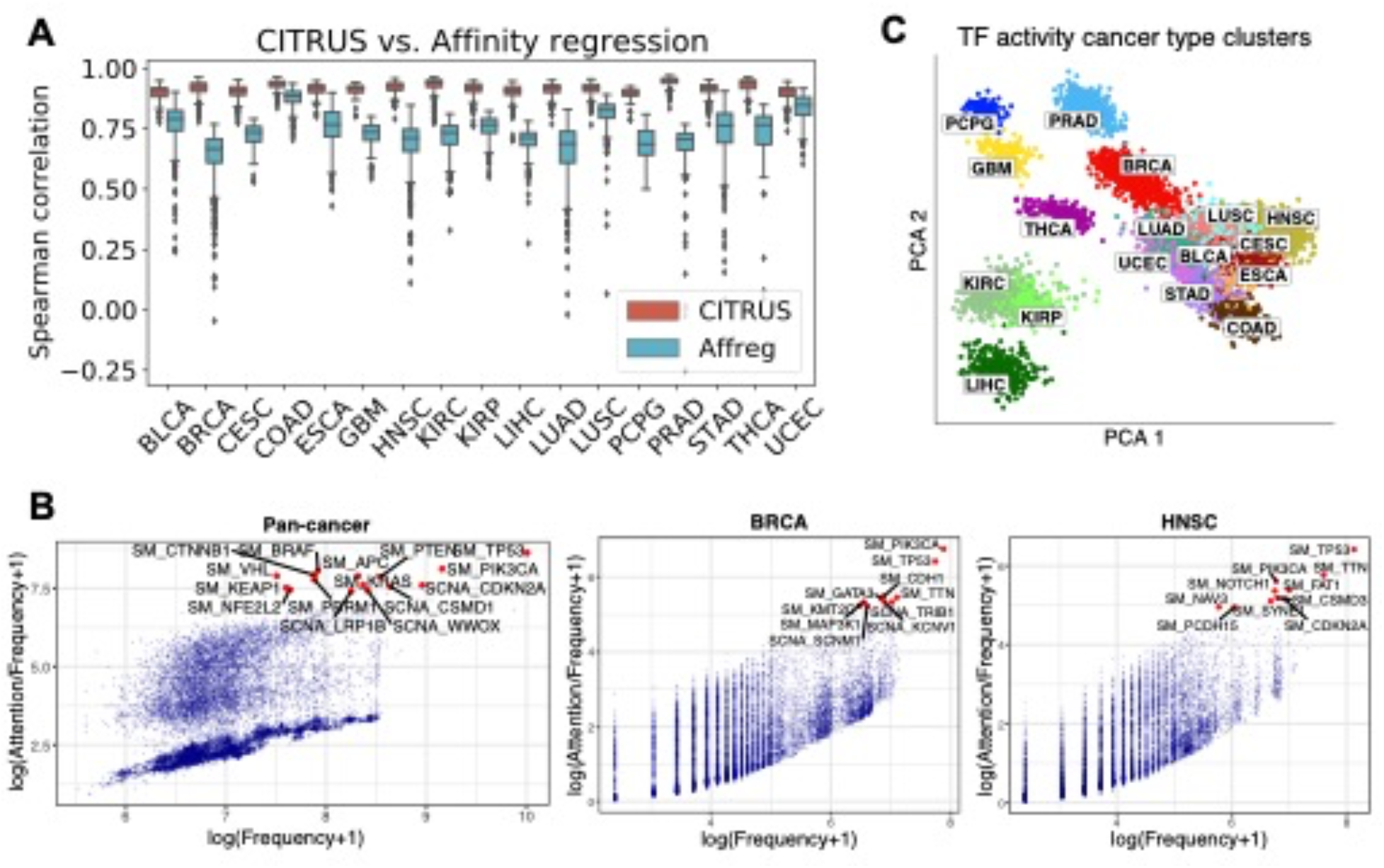
CITRUS models the impact of somatic alterations on gene expression programs. **(A)** Performance of CITRUS in each cancer type compared to the regularized bilinear regression method Affinity regression (Affreg). Boxplots show the mean Spearman correlations between predicted and actual gene expression based on CITRUS (orange) and Affreg (light blue) in TCGA datasets for each cancer type. Both CITRUS and Affreg were tuned on the same training and validation sets and evaluated on the same testing set. **(B)** Somatic alteration frequencies and CITRUS-inferred attention weights of genes. Cumulative pan-cancer results are shown on the left, and individual BRCA and HNSC results are shown in the middle and on the right, respectively. See **Supplementary Fig. 1** for complete results from each cancer type. **(C)** Principal component analysis (PCA) of TF activity colored by cancer type. Standard TCGA tumor symbols are used to indicate tumor type.

To identify somatic alterations that influenced gene expression programs, we compared the relationship of overall attention weights (inferred by CITRUS) and the frequencies of somatic alterations (used as the control group) across all cancer types and within each cancer type (**Fig. 2B and Supplementary Fig. 1**). In general, attention weights were positively correlated with the frequency of somatic alteration. For example, the top altered genes *TP53* and *PIK3CA* had high attention weights. However, our self-attention mechanism assigned low attention weights to many frequently altered genes, indicating that these genes may be cancer passengers. Indeed, we found genes with high attention weights were enriched for known cancer drivers using the IntOGen^9^ database. We first grouped all the genes into two parts with the threshold of 2 (log(attention+1) ≥ 2 as the more attended group, and log(attention+1) < 2 as the less attended group). Using Fisher’s exact test, we verified that known cancer driver genes were enriched in the highly attended group (*P* = 4.48 × 10^−41^) in the pan-cancer analysis. We also observed a few infrequently altered genes with high attention weights. For example, the H3K4 methyltransferase KMT2C had a high attention weight in BRCA but was infrequently altered. Indeed, *KMT2C* is a key regulator of ERα activity and anti-estrogen response in breast cancer (23,24).

We used CITRUS to infer patient-specific TF activities across tumor types. Clustering tumors by these inferred TF activities largely recovered the distinction between major tumor types (**Fig. 2C**). Interestingly, samples with squamous morphology components (BLCA, CESC, ESCA, HNSC, and LUSC) clustered together. Tumors with tissue or organ similarities or proximity were also clustered together. These included neuroendocrine and glioma tumors (GBM and PCPG), clear cell and papillary renal carcinomas (KIRC and KIRP), a gastrointestinal group (COAD, and STAD), and breast and endometrial cancer (BRCA and UCEC). We also observed similar clustering of the tumor embeddings (**Supplementary Fig. 2).**

Next, we assessed TF-tumor type associations by t-test and compared inferred TF activities between samples in each tumor type versus those in all other tumor types. We corrected for false discovery rate (FDR) across TFs and identified significant shared and cancer-specific TFs, which are listed in **Supplementary Data 1**. The average TF activity and significance of the four most significant TFs in each cancer are shown in **Fig. 3**. Our results highlight both known and novel cancer-specific TF regulators. For example, FUBP1, which regulates *c-Myc* gene transcription, had significantly higher inferred activity in many cancer types, including LIHC, HNSC, BLCA, ESCA, CESC, LUSC, PRAD, BRCA, and UCEC. Consistent with previous reports, IRF3 activity was significantly higher in GBM(25). KLF8 had decreased activity in GBM, LIHC, and KIRC, which is consistent with its role in suppressing cell apoptosis during tumor progression (26). Additionally, YY1, which regulates various developmental processes (27), had increased activity in CESC and COAD.

**Fig. 3:**
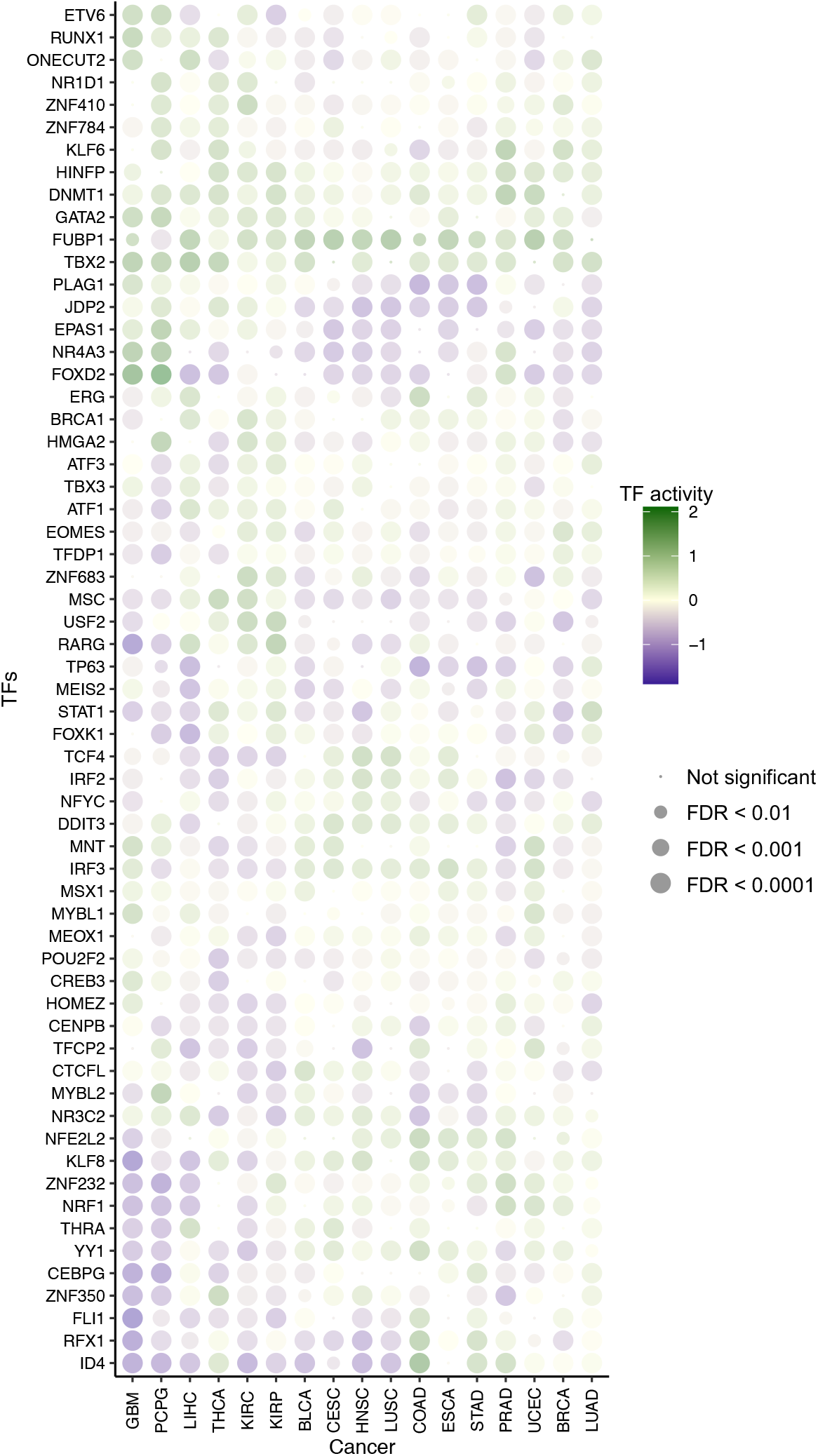
CITRUS identifies regulatory features of tumor types. Dot plot shows the mean inferred TF activity differences between samples in a given tumor type versus those in all other tumor types by t-test. We corrected for FDR across TFs for each pairwise comparison and identified significant TFs. The complete results are included in **Supplementary Data 1**. The dot size indicates −log10(FDR). For clarity, the union of the top four significant TFs in each cancer type is shown.

### Cancer subtype identification from CITRUS-inferred TF activity and somatic alterations

Next, we asked whether CITRUS could identify cancer subtypes based on the TF activity associated with somatic alterations. We conducted *k*-means clustering of inferred TF activities for each cancer type to define subtypes, and then we conducted hierarchical clustering of both the cancer subtypes and TF activities. **Fig. 4** shows the clustering of subtypes by CITRUS-inferred mean TF activities and corresponding somatic alteration associations (see Methods). We observed major differences in mean TF activities across cancer types and minor but significant differences within cancer types. Variations within a cancer type may arise from distinct mutation or CNV profiles of subgroups. For example, clustering by TF activities revealed subclasses of CESC enriched with *KRAS*; KIRC enriched with *VHL*, *BAP1*, *PBRM1,* and *TP53*; LIHC enriched with *CTNNB1*, *BAP1,* and *TP53*; THCA enriched with *NRAS*, *HRAS,* and *BRAF*; and PCPG enriched with *HRAS*.

**Fig. 4:**
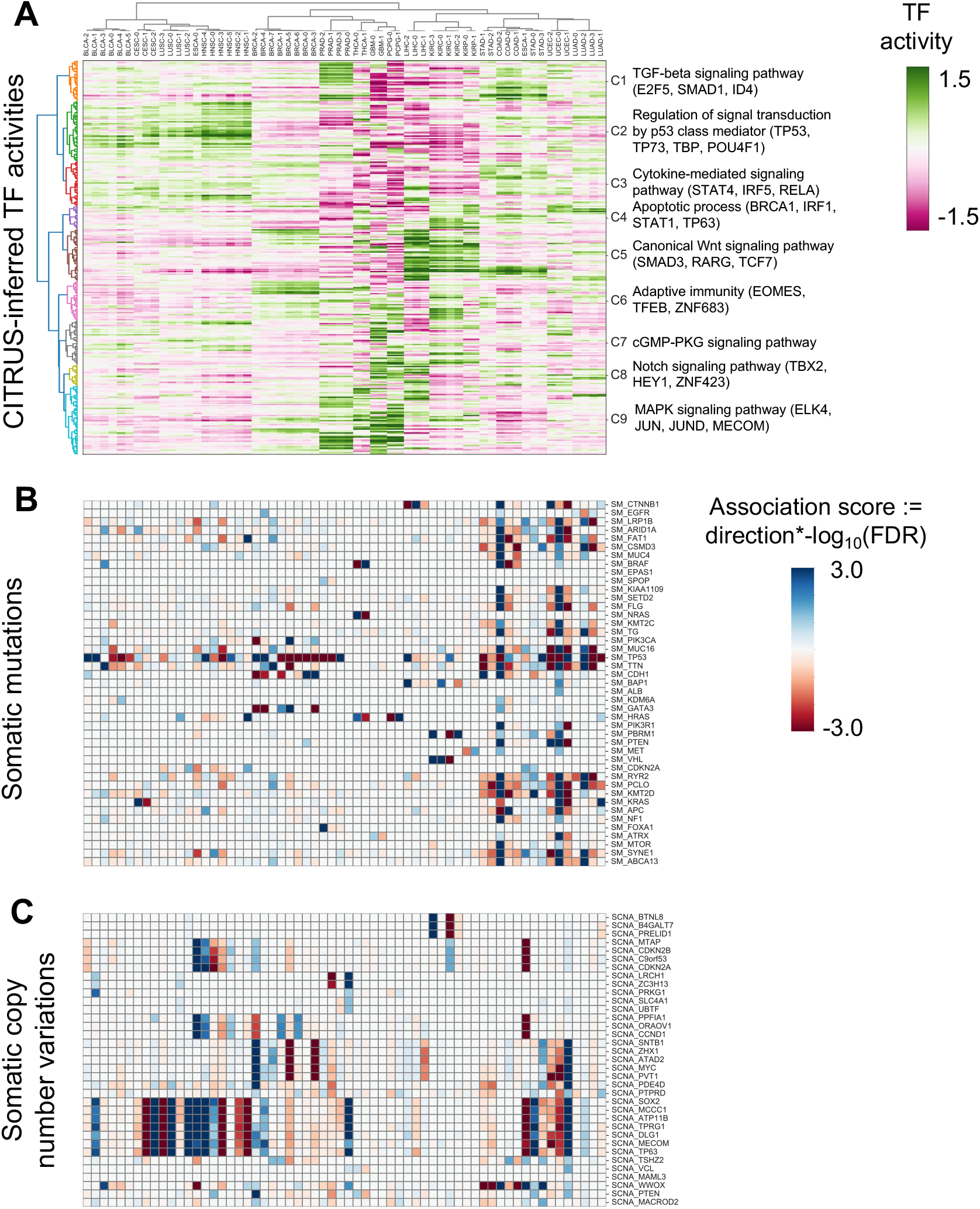
Landscape of somatic alterations and inferred TF activities. **(A)** Heatmap shows tumor subtypes clustered by mean inferred TF activity. The color scale is proportional to TF activity. **(B–C)** Heatmaps of association scores for **(B)** mutations and **(C)** copy number variations. Association scores were calculated by multiplying the −log_10_ FDR by the direction derived from Fisher’s exact test.

As our goal was to decipher cancer-specific downstream effects of targeted therapies and to discover secondary targets for combination drug strategies, we developed a systematic statistical approach for modeling the impact of somatic alterations on TF activity. We implemented an *in silico* knock out approach that removes a specific somatic mutation (or CNV) *g* from all carrier tumor samples in each TCGA cancer study and then predicts altered TF activity (see Methods). Using this approach, we were able to identify TFs whose inferred activity was significantly dysregulated by somatic alterations in known cancer driver genes. **Fig. 5A** demonstrates TF activities that were associated with somatic alterations in UCEC. CITRUS identified mutations in *PIK3CA*, *PTEN*, *KRAS*, *TP53*, and *CTNNB1* that were significantly associated with various TF activities across UCEC tumors (~66% of tumors have *PTEN* inactivating mutations, ~50% have *PIK3CA* activating mutations, ~38% have *TP53* mutations, ~26% have *CTNNB1* mutations, and ~20% have *KRAS* mutations). UCEC samples with *PTEN* mutations were mutually exclusive with *TP53*, *CTNNB1,* and *KRAS* mutations and showed distinct TF activity patterns. Mutations in *PTEN* that inactivate its phosphatase activity result in increased PI3K signaling. Consistent with this effect, TFs associated with *PTEN* mutations were involved in cell cycle and differentiation, including E2F5, TP63, ELF3, DBP, ZKSCAN3, LHX2, HOXB6, SOX9, DBP, MYLB1, and GLIS1. TFs associated with *CTNNB1* mutant status were involved in WNT and TGF-beta signaling including TCF7, TCF7L2, TCF7L1, FOXH1, EMX1, and MYBL1.

**Fig. 5:**
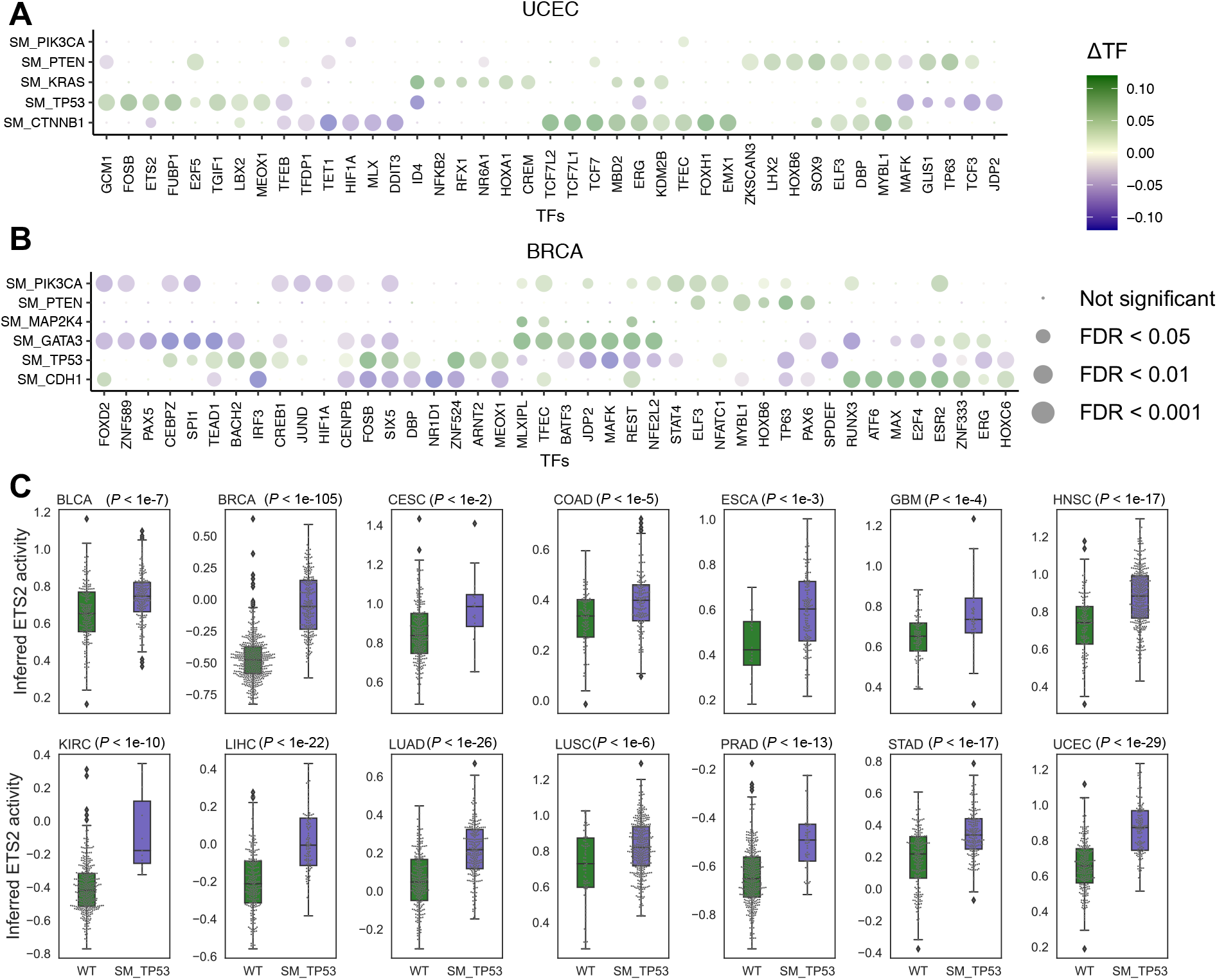
Somatic alterations are associated with dysregulated TF activity. Impact of somatic alterations on individual TFs based on *in silico* knock out experiments in **(A)** UCEC and **(B)** BRCA datasets from TCGA. The dot plot shows mean TF activity, and dot size indicates −log10(FDR). See **Supplementary Fig. 3** for the full list of cancer types. **(C)** Inferred ETS2 activity in TCGA studies and impact of *TP53* mutations. Tumors with mutant *TP53* have significantly higher ETS2 activity than WT tumors (*P* < 0.01, t-test). This association is not significant using mRNA levels of *ETS2* (**Supplementary Fig. 5)**. Box edges represent the upper and lower quantile with median value shown as a bold line in the middle of the box. Whiskers extend to 1.5 times the quantile.

Similarly, CITRUS identified TF activities that were associated with somatic alterations in BRCA (**Fig. 5B**). Mutations in *PIK3CA*, *PTEN*, *MAP2K4*, *GATA3*, *TP53*, and *CDH1* were significantly associated with various TF activities. In BRCA, ~36% of tumors have *PIK3CA* activating mutations, ~35% have *TP53* mutations, ~15% have *GATA3* mutations, ~15% have *CDH1* mutations, ~10% have *PTEN* mutations, and ~7% have *MAP2K4* mutations. Activating mutations in *PIK3CA* often occur in one of three hotspot locations (E545K, E542K, and H1047R) and promote constitutive signaling through the pathway. TFs associated with *PIK3CA* mutations were involved in WNT signaling, epithelial–mesenchymal transition, and cancer stem cell transition, including ELF3, TFEC, STAT4, STAT5B, NFATC1, GLIS1, CDC5L, and AR. BRCA samples with *PIK3CA* and *TP53* mutations were mutually exclusive, and our *in silico* knock out analysis associated distinct TFs with these mutations. *TP53* mutant tumors were associated with increased activity of TFs that have roles in tumor growth, such as ETS2 and FOSB, growth modulation, such as THAP1, CREB3L1, and CEBPZ, and development, such as MEF2C/D, MEOX1, and MSX1. We performed similar analyses for other cancer types (**Supplementary Fig. 3**).

Although the TFs affected by some somatic alterations differed between cancer types, mutation of *TP53* was associated with similar TFs across cancer types (**Supplementary Fig. 4**). *TP53* is one of the most frequently inactivated tumor suppressor genes that suffers from missense mutations in human cancer. These missense mutations result in the expression of a mutant form of p53 protein. Mutant p53 protein can disable other tumor suppressors (e.g., p63 and p73) or enable oncogenes, such as ETS2 (28). Indeed, the inferred TF activity of ETS2 was higher in mutant versus WT *TP53* tumors across cancers (**Fig. 5C**); however, these differences were not as significant at the gene expression level (**Supplementary Fig. 5**).

## Discussion

Analysis of the regulatory network in tumor datasets is challenging due to the complexity of the cancer genome (e.g., aneuploidy, CNVs, structural variation, and mutations). CITRUS provides a systematic framework for integrating regulatory genomics with tumor expression and somatic alterations to better understand how expression programs are affected by somatic alterations in cancers and to infer patient-specific TF activities. Our method uses a deep learning framework called a self-attention mechanism to capture the complex contextual interactions between somatic alterations. For a more accurate representation of TF-target gene relationships, we leveraged ATAC-seq tumor data from TCGA patients. CITRUS is designed to capture the flow of information from altered genes (e.g., signaling proteins) to TFs to target genes, and our *in silico* knock out analysis predicts the causal impacts of somatic alterations. Joint modeling across different tumor types also revealed patient subgroups associated with somatic alterations. In cases where a somatic alteration is associated with the activity of a targetable TF or their upstream/downstream component, it may be possible to identify combination therapies using CITRUS.

One limitation of the TF binding motif approach utilized by CITRUS is that TFs of the same family often share a similar motif and thus are difficult to disambiguate. Therefore, TF motifs may encompass the activities of multiple TFs. Moreover, co-binding TF binding patterns (e.g., AP-1−IRF complexes) can be biologically meaningful for gene expression and are not currently represented in our model. Future models will work to represent these composite elements as features. Another limitation is that we do not represent directionality in the TF-target gene priors (i.e., whether a gene is activated or repressed by a TF). Prior knowledge of whether the TF is acting as an activator or as a repressor would add meaningful interpretation to inferred TF activities. These limitations may confound the interpretation of the activity of TFs with context-specific activator and repressor roles. Further, regulatory network analysis of tumor datasets is also complicated by the presence of stromal/immune cells within the tumor and the heterogeneity of the cancer cells themselves. However, our framework can be extended to model single-cell RNA-seq or deconvoluted RNA-seq via computational methods.

Despite these limitations, modeling the impact of somatic alterations on transcriptional programs may ultimately enable the development of individualized therapies, aid in understanding mechanisms of drug resistance, and allow the identification of biomarkers of response. We anticipate that computational modeling of transcriptional regulation across different tumor types will emerge as an important tool in precision oncology, aiding in the eventual goal of selecting the best therapeutic option for individual patients.

### Data availability

ATAC-seq data are available in the public repository Genomic Data Commons (https://gdc.cancer.gov/about-data/publications/ATACseq-AWG). RNA-seq gene expression, somatic mutation, copy number variation, and clinical data are available in a public repository from TCGA’s Firehose data run (https://confluence.broadinstitute.org/display/GDAC/Dashboard-Stddata). Only the samples ‘whitelisted’ by TCGA for the Pan-Cancer Analysis Working Group were used in the study. For our analysis, we only used samples with parallel RNA-seq, somatic mutation, and GISTIC copy number data. Processed input and output files have been made available at the supplementary website for the paper: http://www.pitt.edu/~xim33/CITRUS.

### Code availability

The software for CITRUS is available at https://github.com/osmanbeyoglulab/CITRUS.

## Acknowledgements

The results published here are based on data generated by The Cancer Genome Atlas project established by the NCI and NHGRI (accession number: phs000178.v7p6). Information about TCGA and the investigators and institutions that constitute TCGA research network can be found at http://cancergenome.nih.gov/. We thank Jacob Stewart-Ornstein for helpful discussions

